# Integrated deep learned transcriptomic and structure-based predictor of clinical trials outcomes

**DOI:** 10.1101/095653

**Authors:** Artem V. Artemov, Evgeny Putin, Quentin Vanhaelen, Alexander Aliper, Ivan V. Ozerov, Alex Zhavoronkov

**Affiliations:** Pharmaceutical Artificial Intelligence Division, Insilico Medicine, Inc., Emerging Technology Centers, Johns Hopkins University at Eastern, B301, 1101 33rd Street, Baltimore, Maryland 21218, USA

**Keywords:** Clinical trials, Machine learning, Deep learning, Transcriptome, Side effects

## Abstract

Despite many recent advances in systems biology and a marked increase in the availability of high-throughput biological data, the productivity of research and development in the pharmaceutical industry is on the decline. This is primarily due to clinical trial failure rates reaching up to 95% in oncology and other disease areas. We have developed a comprehensive analytical and computational pipeline utilizing deep learning techniques and novel systems biology analytical tools to predict the outcomes of phase I/II clinical trials. The pipeline predicts the side effects of a drug using deep neural networks and estimates drug-induced pathway activation. It then uses the predicted side effect probabilities and pathway activation scores as an input to train a classifier which predicts clinical trial outcomes. This classifier was trained on 577 transcriptomic datasets and has achieved a cross-validated accuracy of 0.83. When compared to a direct gene-based classifier, our multi-stage approach dramatically improves the accuracy of the predictions. The classifier was applied to a set of compounds currently present in the pipelines of several major pharmaceutical companies to highlight potential risks in their portfolios and estimate the fraction of clinical trials that were likely to fail in phase I and II.

## Introduction

Throughout the last decade many studies have revealed a continuous increase in the total cost of developing successful pharmaceutical products (Munos and Chin 2011; Mignani et al. 2016; Scannell et al. 2012). The estimated drug development costs exceed US$2.5 billion per the new molecular entity (NME) and the returns of investment in breakthrough drugs with the exception of cancer immunology, are on the decline (Munos and Chin 2011; Mignani et al. 2016; Rubin and Gilliland 2012; DiMasi, Grabowski, and Hansen 2016). This forces many pharmaceutical companies to opt for not taking on ambitious innovation plans that might have resulted in a range of investigational drugs for addressing various health issues faced by both developed and developing countries around the world (Hay et al. 2014; Ledford 2011). The rising cost of in vivo studies dominating the total development cost is a major concern, especially taking into account the very low predictive value for human clinical trials (Viceconti, Henney, and Morley-Fletcher 2016). Despite the many efforts and substantial resources invested in the identification and careful design of new compounds and the systematic monitoring of all steps in the early development process and extensive in vitro validation, even the most promising high-value compounds often fail in clinical trials. Improvements in the drug development process failed to resolve the disturbingly high and continuously growing failure rates in clinical trials due to issues with safety and efficacy (Hay et al. 2014; Ledford 2011). In 2012, only 39 drugs classified as NMEs and biologic license applications (BLAs) were approved by the FDA (Mullard 2013). The success rates vary among different disease areas. In oncology only 1 of 15 drugs that enter phase I clinical trials are approved by the FDA (Hay et al. 2014; Rubin and Gilliland 2012), and for cardiac medications, where none of prospective compounds from phase I and II studies have been successful in phase III clinical trials in recent years (Lavine and Mann 2013). Multiple specific solutions have been proposed to address the common causes of failure at each phase of clinical trials (Berry 2011). Finding innovative solutions to reduce the clinical failure rates is one of the main challenges of the pharmaceutical industry as a failures during late-stage clinical trials result in substantial losses. The fact that clinical trials alone do not provide enough information about the structural and design properties explaining a candidate’s failure makes a repurposing process of such compound difficult(Viceconti, Henney, and Morley-Fletcher 2016). Often times this leads to its complete abandonment at the expense of considerable resources invested in its early development, despite the fact that such compounds could have been repositioned as candidates for other conditions (Ashburn and Thor 2004; Vanhaelen et al. 2016).

Although many factors contribute to this concerning trend, it is acknowledged that significant drug discovery workflow optimization is necessary in order to fully integrate the knowledge about the onset and progression of diseases gained over the last decades with the latest technological and computational advances. The standard workflow employed by pharmaceutical companies for developing a new compound can be roughly divided into three main phases: design/discovery, pre-clinical assessment and clinical assessment (Viceconti, Henney, and Morley-Fletcher 2016). Currently it is common practice to use various computational modeling methods and simulation tools to accelerate and improve the efficiency of drug discovery, design and other steps in the early development stage. The fact that over 75% of the development cost of a new compound is spent on in vivo studies (Viceconti, Henney, and Morley-Fletcher 2016) implies that improvements made at the level of the early development in silico cycle have a small impact on the total development cost but could significantly help with predicting the outcome of clinical trials. Furthermore, integrating in silico approaches inside the late development phases and making use of in silico clinical trials (ISCT) could significantly reduce the cost of clinical trials while improving the overall success rate (Chabanas, Luboz, and Payan 2003; Fernandez and Hunter 2005; Li et al. 2008; Clermont et al. 2004). Currently, one of the most ambitious projects in this direction is the initiative called Virtual Physiological Human (VPH) (Viceconti, Henney, and Morley-Fletcher 2016). The final goal of this project is to develop a framework where various types of technologies could be used to generate quantitative information about the biology, physiology, and pathology of a patient at different scales of space and time, with the ultimate aim of producing patient-specific predictions for diagnosis, prognosis, and treatment planning.

Although very attractive, the idea of using ISCT for reducing, refining or partially replacing in vivo experimentation faces three main challenges. Firstly, pharmaceutical companies still lack the fundamental and technological knowledge required to do so, and one has to continuously address the inherent complexity associated with the accurate, quantitative modelling of living organisms. Secondly, the main industry players still need to be convinced of the reliability of in silico approaches, especially in the case of predictions made in place of in vivo experiments. Finally, the adoption of such disruptive technologies can be a tedious process involving changes in business practice and adaptation of the regulations regarding the standard certification process of new pharmaceutical compounds (Viceconti, Henney, and Morley-Fletcher 2016).

Taking these aspects into account, other paths for improving the success rates of clinical trials have been investigated. Another computational-based strategy currently followed is to develop a set of criteria that could be used to predict the outcome of a clinical trial before it actually begins. A major focus of such approaches lies in the ability to identify compounds with unfavorable toxicity properties. Within the pharmaceutical industry, drug-likeness measures are commonly used as guidelines for filtering out toxic molecules in the early development stages. The procedure is based on a set of physical and chemical features associated with orally active drugs obtained from the screening of clinical drugs that reached phase II trials or beyond (Lipinski et al. 2001). This procedure contributed to improving the drug discovery process by providing a set of practical filters for many drug development pipelines (Leeson and Springthorpe 2007). Nevertheless, the initial drug-likeness approach as well as the improved versions developed later (Veber et al. 2002; Ghose, Viswanadhan, and Wendoloski 1999) are not able to distinguish unacceptable toxicity profiles from safe ones (Bickerton et al. 2012; Leeson and Springthorpe 2007). Further modifications to these protocols primarily focus on properties associated with bioavailability, ignoring other aspects that can trigger clinical trial toxicity events.Various computational methods have been developed for tackling the issue of toxicity in various situations, including tissue-specific toxicities (Jeliazkova and Jeliazkov 2011) and QSAR methods used for predicting specific toxicity endpoints and estimating maximum tolerated dose levels (Patlewicz and Fitzpatrick 2016). One recently developed approach called PrOCTOR (Gayvert, Madhukar, and Elemento 2016) takes into account not only bioavailability related properties, but also target-related properties, such as tissue selectivity and established informative chemical features of the drugs with target-based features to produce a classifier based on random forests that can be used to address clinical drug toxicity. In their study, the authors demonstrated that some of these properties, even when taken into account individually, have a significant discriminative power. Furthermore, using a feature importance analysis, they have shown that both the chemical and target-based features contribute significantly to an efficient classification.

One of the emerging fields in machine learning is deep learning, a set of techniques aimed to learn a high level of feature representation. On large datasets in image, text and voice recognition deep learning systems drastically outperform the conventional methods and achieve human-competitive levels of accuracy (LeCun, Bengio, and Hinton 2015). However, there is a rapidly growing amount of literature on the application of deep learning techniques to biomedicine, which indicates that deep learning actively dominates the field of computational biology with its sparse and often limited data. Deep-learned systems have demonstrated reasonable accuracy when guessing patients’ chronological age using anonymized blood biochemistry samples (Putin et al. 2016). The ensemble of neural networks with different architecture and set of hyperparameters outperformed baseline machine learning technique and proved the previously questionable utility of basic blood biochemistry data in precise age prediction. Such spectacular predictive power of neural networks, however, like with any other method, clouded by the fact that population-specific bias could decrease the accuracy of the predictor (Cohen et al. 2016). However, with an increasing number of population-specific samples deep learning methods overcome such training limits in most cases (Mamoshina et al. 2016). Multi-layer neural networks were able to classify small molecules by pharmaceutical classes using one the most complex and heterogeneous categories of biological data - namely, gene expression (Aliper et al. 2016). In order to overcome the “curse of dimensionality” networks were trained on both a set of landmark genes as well as signaling pathways. Pathway-based deep neural networks show better classification accuracy and could be considered as powerful repurposing tools. Another direction in deep learning holding a lot of promise for drug discovery and biomarker development are the Generative Adversarial Networks (GANs). GAN is a fresh direction in deep learning invented by Ian Goodfellow in 2014 (Goodfellow et al. 2014). In recent years GANs produced extraordinary results in generating meaningful images according to the desired descriptions. Similar principles can be applied to drug discovery and biomarker development. The proof of concept of a GAN-based drug discovery engine was developed using the generative adversarial autoencoders (AAEs), (Makhzani et al. 2015) to generate molecular fingerprints of prospective compounds with the desired molecular properties (Kadurin et al. 2016).

Insilico Medicine developed a comprehensive and constantly evolving drug discovery pipeline utilizing deep learning (Figure 1). In this study, we have thoroughly analyzed transcriptomic data for drug-induced perturbations in cell cultures. We applied various machine learning and pathway analysis techniques to predict the side effects of drugs and the consequent success or failure of clinical trials. We demonstrated how pathway analysis and the accurate prediction of side effects can help to achieve high predictive accuracy of clinical trial outcomes. Finally, we applied the classifiers constructed on retrospective data to the current pipelines of the major pharmaceutical companies.

**Figure 1.**
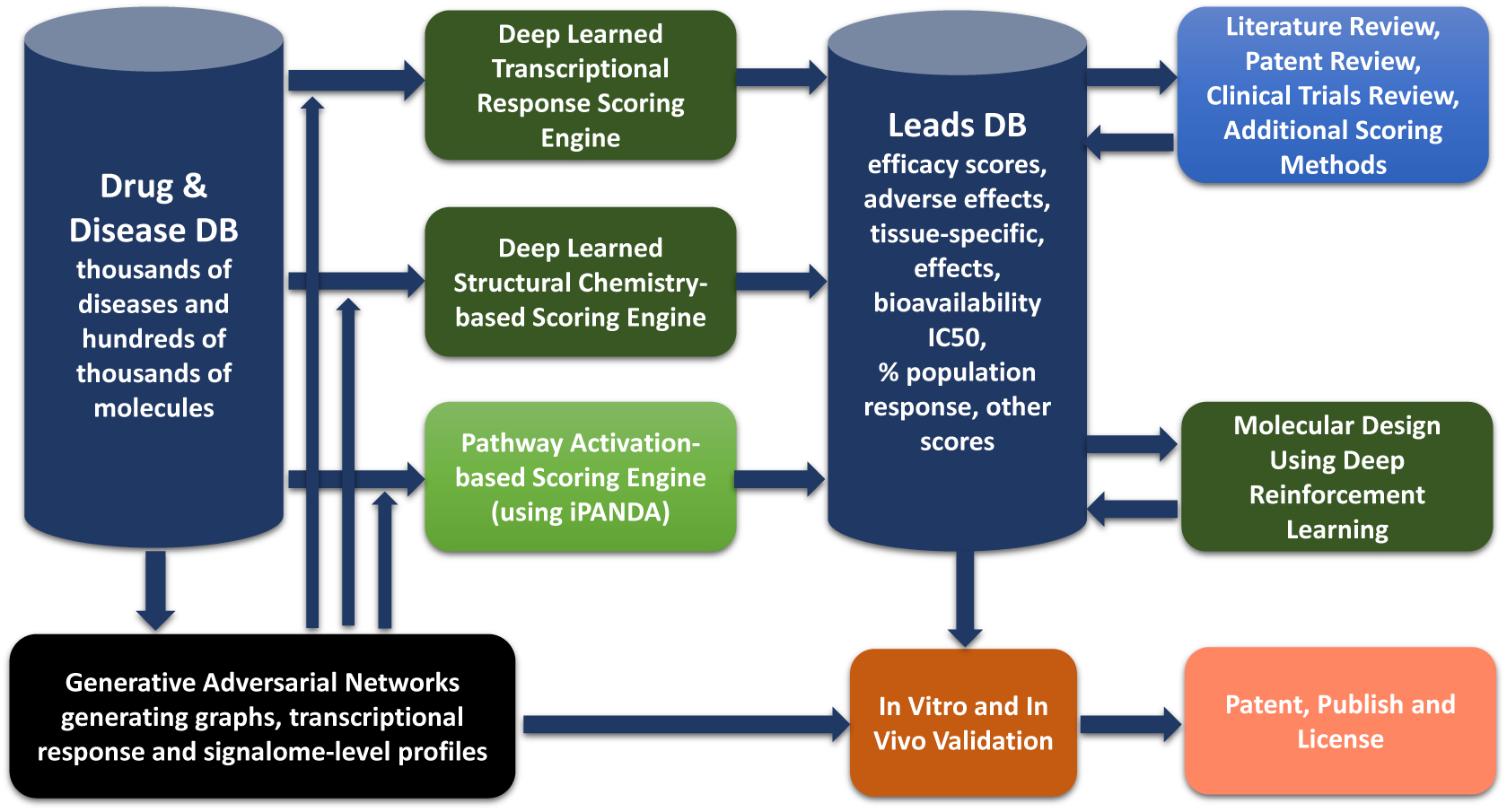
Artificially intelligent pipeline for generating drug leads. Insilico Medicine developed a drug discovery pipeline utilizing a large number of data sets of multiple data types. The pipeline starts with a database of transcriptional response and structural data linked to other molecular properties that is supplied into the transcriptomic drug discovery engine, structural drug discovery engine and pathway-centric drug discovery engine. The data is also used to train the generative adversarial networks to provide leads in the form of graph representations of molecular structure that can be supplied into the structure-based predictor, or tested experimentally. It also generates transcriptional response data sets for pathway-based and deep learned predictors. After the leads are ranked by the efficacy, safety and other properties, the literature and patent review is performed. For molecules that can not be patented, another engine utilizing deep reinforcement learning is used to develop analogs that may have the same or superior properties and can be tested and patented.

## Results

### Neural networks accurately predict common side effects

Deep neural networks were constructed to predict side effects of compounds based on the transcriptional alterations they cause in cell lines. As an input, the neural networks took pathway activation scores, estimated by the iPANDA algorithm (Ozerov et al. 2016). These scores represented activation or inhibition of various regulatory pathways following treatment of a cell line with a given compound. We built a single neural network for each side effect. The accuracy, sensitivity, specificity and f1 measure were estimated after cross-validation for each network. For 46 side effects, we achieved f1 greater than 0.8.

### Gene expression changes after drug treatment poorly predict clinical trials outcome

We firstly investigated whether the drug-induced changes of gene expression *per se* could predict the outcome of a clinical trial. We constructed a set of classifiers based on relative gene expression changes following treatment with a compound of interest. The random forest (RF) classifier achieved an accuracy of 0.66 (Figure 2B).

**Figure 2.**
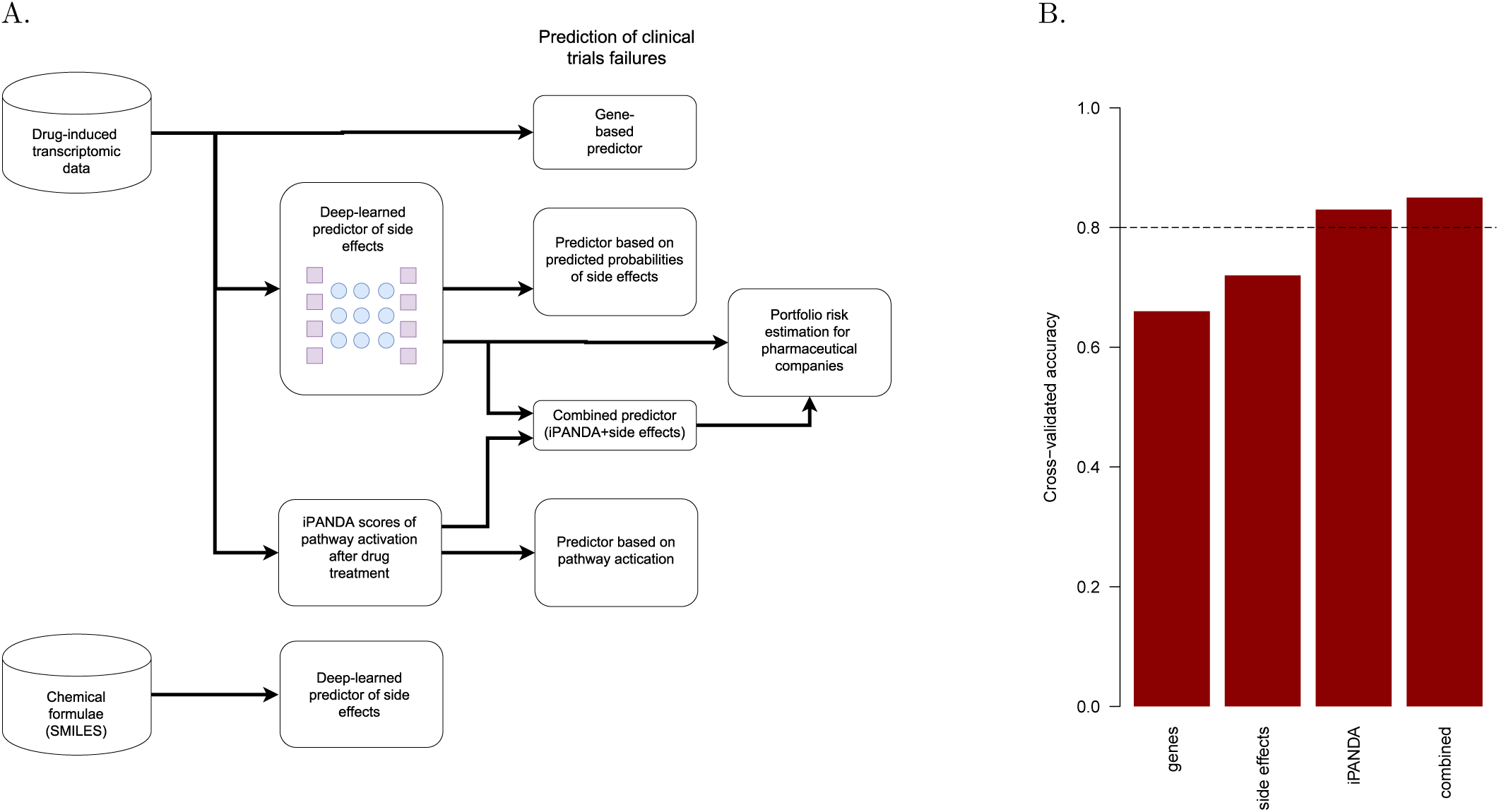
**(A)** A schematic representation of the computational experiment. **(B)** Cross-validated accuracy of clinical trials’ outcome prediction for the following classifiers: random forest based on drug-induced gene expression changes; random forest based on probabilities of various side effects predicted by a deep neural network; random forest based on iPANDA pathway activation scores after drug treatment; combined random forest classifier utilizing both side effects predictions and iPANDA scores.

### Deep-learned predictions of side effects explain failures of Phase II

Unexpected side effects could be one of the reasons for clinical trial failure. Therefore, we explored whether our transcriptomic-based predictions of side effects could be used to predict the outcome of clinical trials in Phase II. We first selected the side effects for which we had high confidence of the prediction given by a neural network, namely those with f1 measure greater than 0.8. Next, we built a random forest classifier based on probabilities of each side effect to be discovered for each compound, predicted by the neural network. The classifier achieved much higher accuracy (0.81) compared to analyses based on gene expression changes as is (Figure 2B).

### Pathway-based classifier predicts the outcome of clinical trials

We hypothesized that the low accuracy of the gene-based classifier could be explained by the high number of features (genes with measured expression) compared to the number of samples (i.e., compounds for which the outcomes of clinical trials are known). Therefore, an efficient and biologically relevant method for dimensionality reduction could potentially improve the accuracy. To test this hypothesis, we applied the iPANDA algorithm (Ozerov et al. 2016) to estimate the activation or inhibition of certain regulatory and biochemical pathways following treatment with a drug of interest. The iPANDA scores were then used as an input for the classifier designed to predict the outcomes of clinical trials (Figure 2A). The obtained results show that using pathway activation scores as predictors in the classifier dramatically improved its accuracy up to 0.83 (Figure 2B).

In order to understand which pathway activation had the greatest contribution to the prediction, we performed importance ranking of the input variables. In Supplementary Figure S1, we summarized the most important variables for the classifier. A significant number of metabolic pathways, in particular those involved in oxidation, as well as endoplasmic reticulum (ER)—related pathways, were among the best predictors of clinical trial failures. Interestingly, these processes are responsible for metabolism of xenobiotics and are particularly active in hepatic tissue.

### A combined classifier accounts for both side effects and efficacy

To account for both potential side effects and the mode of action of each compound of interest, we constructed a combined classifier which used both the predicted probabilities of side effects and iPANDA pathway activation scores. This combined model showed performance comparable to the model based on iPANDA scores alone (Figure 2B) and therefore we used the classifier based on iPANDA scores for the further analysis.

### Analysis of pharma pipelines

The classifier achieved a cross-validated accuracy of 0.83, which is high enough to predict the outcome of an individual clinical trial. It was even more suitable to assess the portfolio of compounds to estimate the fraction of compounds under the risk of failing a clinical trial, as the uncertainty in the prediction for an individual compound is compensated by the number of compounds in a portfolio. We predicted probabilities of failure in clinical trials for the compounds currently included in the pipelines of one of four major pharmaceutical companies (Figure 3A–D).

**Figure 3.**
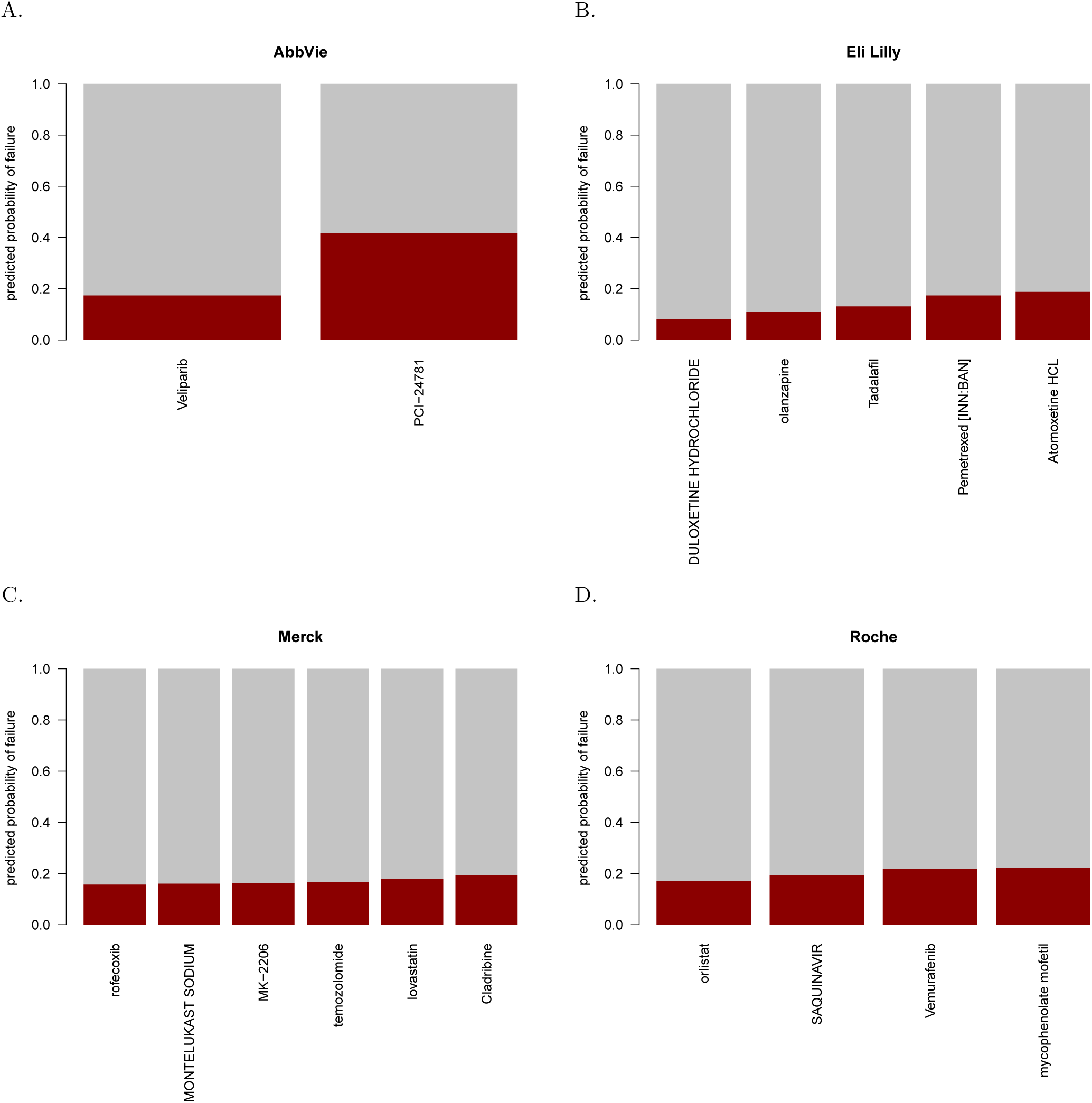
Analysis of drug pipelines for certain major pharmaceutical companies. Red bars show the predicted probabilities of clinical trials’ failures for individual compounds. **(A)** AbbVie, **(B)** Eli Lilly, **(C)** Merck, **(D)** Roche

In our prediction, Abexinostat (PCI-24781), an HDAC inhibitor developed by AbbVie, had a high expected probability of failure. There are several completed clinical trials featuring this compound. First Phase I trial (NCT01149668) was focused on investigating long-term safety of Abexinostat in subjects with various hematologic cancers. Although the trial was completed in 2013, the results have not been published so far. Second trial Phase I/II trial (NCT00724984) had a slightly different scope and focused only on patients with Hodgkin’s and Non-Hodgkin’s lymphoma. Recently published results (Evens et al. 2016) indicate that the drug was well tolerated and showed good efficacy in follicular lymphoma. Grade 3-4 treatment-related adverse events in Phase II were thrombocytopenia, fatigue, and neutropenia.

For an independent evaluation of our pipeline’s ability to detect successes (Figure 4A) and failures (Figure 4B) of clinical trials, we applied the classifier to a set of novel compounds in the new L1000 Connectivity Map perturbational profiles dataset (GSE70138). We predicted the highest probabilities of failure for PKC inhibitor enzastaurin, CDK inhibitor dinaciclib (MK-7965), prospective Hsp90 inhibitor tanespimycin, BET bromodomain inhibitors I-BET151 (GSK1210151A) and JQ1, PKC/CDK inhibitor CGP60474 and B-Raf inhibitor SB590885 (Figure 4B). None of these compounds were approved: according to Lilly report, enzastaurin failed in Phase III; the clinical trials of tanespimycin were halted by Bristol-Myers Squibb; dinaciclib (MK-7965) showed severe side effects and no improvement of efficacy in non-small cell lung cancer and breast cancer clinical trials (Stephenson et al. 2014; Mita et al. 2014). In contrast, three among eight of compounds for which we predicted the lowest probabilities of failure, namely entinostat, vorinostat and resveratrol, did in fact pass phase II clinical trials or were approved by FDA (Figure 4A).

**Figure 4.**
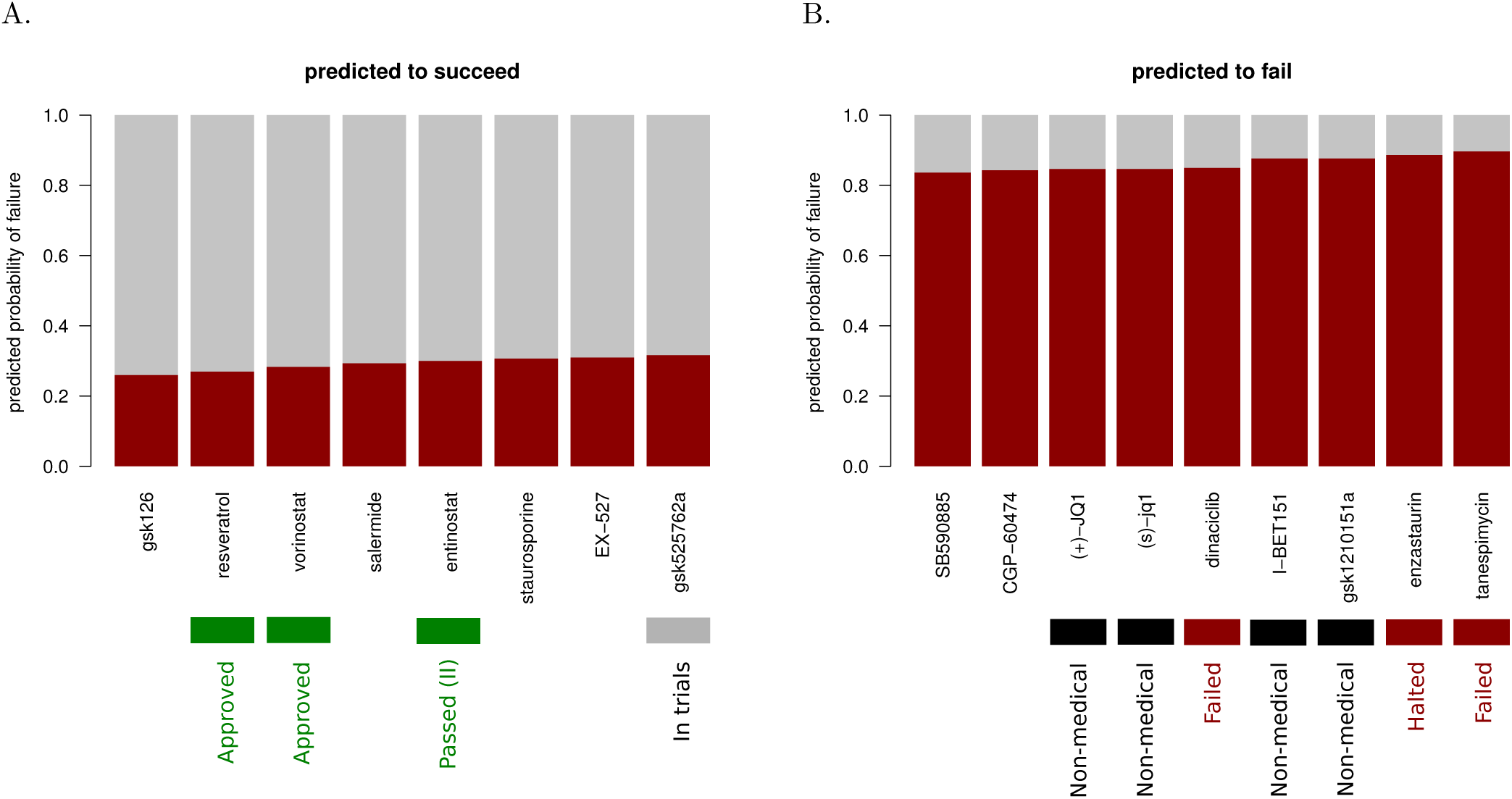
Compounds predicted to succeed and fail in clinical trials from an independent transcriptomic dataset. Red bars show the predicted probabilities of clinical trials’ failures for individual compounds. Available clinical trials data is summarized in the bottom. The compounds which are not intended to be used medically are marked as non-medical. Panel **(A)** depicts the compounds with the lowest predicted probabilities of failure while panel **(B)** depicts the compounds with the highest predicted probabilities of failure.

## Methods

### Data preparation

As an input, we used the LINCS L1000 dataset with transcriptional response of multiple cell lines to drug treatment. In the dataset, expression was profiled for a thousand “milestone” genes (Lamb et al. 2006). For a naïve gene expression-based classifier, we performed limma test of differential gene expression between the samples treated with a compound and control samples of a given cell line. The obtained signed values of test statistics were used as an input for the gene-based classifiers.

For pathway-based classification, we calculated pathway activation scores using the iPANDA algorithm (Ozerov et al. 2016). We used a collection of signalling pathways from KEGG (Kanehisa 2000), NCI (Schaefer et al. 2009) and SABiosciences (http://www.sabiosciences.com/pathwaycentral.php) pathway databases.

Clinical trials data was processed similarly to the approach used in (Gayvert, Madhukar, and Elemento 2016).

### Deep neural network for side effects prediction based on transcriptomic data

Deep neural networks (DNNs) were trained with transcriptional response data from the LINCS L1000 dataset. Side effects for drugs were derived from SIDER database (Kuhn et al. 2015). Side effect categories were mapped onto 205 preferred terms from MedDRA v16.0 ontology (Brown, Wood, and Wood 1999). Each DNN was trained to predict concrete side effect, thus each DNN was binary. In total 205 DNNs were trained. DNNs were trained with Dropout (Srivastava et al. 2014) and l2 regularization to prevent overfitting. For each side effect the optimal topology (number of neurons in each layer, number of layers, optimizer, etc) of the DNN in terms of accuracy was found by grid search procedure. The search space for all DNNs was: from 100 to 1000 neurons, from 3 to 6 layers, Adam (Kingma and Ba 2014), Adadelta (Zeiler 2012), Adagrad (Duchi, Hazan, and Singer 2011) as optimizers, ReLU, PReLU, ELU as activation functions, from 0.25 to 0.5 as dropout probability. For all DNNs binary cross entropy was used as a loss function. DNNs were trained on NVIDIA Titan X graphical processing unit with 5-fold cross validation.

### Classification and cross-validation

The classification based on RF was performed using the randomForest function from the standard R distribution and the glm function (with parameter method=”binomial”) was used for logistic regression. 500 trees were used to construct a random forest classification.

The classifier was trained on 577 transcriptomic datasets obtained for 214 compounds.

Multiple lines of the input data matrix could correspond to a single drug (e.g., a drug can be profiled at different concentrations, cell lines and durations of treatment). To prevent the data related to a single drug from appearing in both the training and the test set, the following cross-validation procedure was applied: we extracted the list of unique drug identifiers and on each iteration took a random 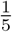 of those as a test set of compounds; only the lines of the input data matrix not corresponding to the drugs in the test set were considered for training.

Before predicting probabilities of failure for the compounds of a certain company, we excluded the compounds of that company from the training set and rebuilt the classifier. The classifier (trained on all compounds except the compounds of the analyzed company) was used to predict probabilities of failure for the compounds of the given company.

## Discussion

In this work, we described certain computational methods to predict drug efficacy and clinical trial failure or success based on transcriptomic data. As expected, the outcome of a clinical trial appeared to depend both on potential side effects and on the efficacy of the drug. This can explain the fact that the combined classifier performed better than those based on either gene expression or predicted side effects alone.

Both the iPANDA-based classifier and the combined classifier achieved high cross-validated performance. Remarkably, the results of iPANDA-based DNNs for side effects significantly outperform gene-based DNNs. Previously we showed that DNN classifiers of therapeutic category trained on signaling pathway activation scores are significantly more accurate than their gene-based counterparts (Aliper et al. 2016).

Overall, such prediction tools can be of a great value, especially at the stage of lead screening. The only type of data required to perform the prediction is gene expression before and after incubation of a single cell line with the compound of interest, which can be obtained in a cost effective and high-throughput manner.

We aimed to analyze the current pipelines of the major pharmaceutical companies to estimate what proportion of currently tested drugs is likely to fail Phase 2 clinical trials. This estimation can be thought of as a clinical analog of the value-at-risk concept. We applied the classifiers to the current pipelines of pharma companies and obtained the risk rate, or clinical trial failure rate, as well as predictions of multiple side effects.

Rather low predicted probabilities of failure for the analyzed compounds could seem contradictory to the overall 70% failure rate. This contradiction can be resolved by taking into account the fact that the compounds for which L1000 transcriptomic data is available represent a biased set of compounds: they are on the market for a longer time than the average compound, and some of them have already been successfully tested for other indications. Therefore, the analyzed set of drugs can be less likely to fail than the average drug in the current R&D pipelines.

In contrast, among the novel compounds in the new L1000 dataset, which are currently studied in preclinical and clinical trials, we discovered those which were likely to fail (Figure 4B). Most of our predictions already failed the clinical trials due to severe side effects or lack of improvement in efficacy. Interestingly, some BET inhibitors showed high probabilities of failure in our analysis, which corresponded to their observed side effects. Moreover, while we predicted failure for BET inhibitors JQ-1 and I-BET151 which were not intended to be used medically, we predicted success for their analog gsk525762a (I-BET 762) which is currently in clinical trials (Figure 4A). This advocates for a more thorough evaluation of candidate BET inhibitors in preclinical studies. As BET inhibitors have a direct and broad effect on gene expression, our transcriptomic-based pipeline can be particularly useful for the prediction of side effects and clinical trial failures for the members of this class of compounds.

Interestingly, the studied companies showed different risk rates. Among four investigated big pharma companies, AbbVie has the highest median risk rate. On the other hand Eli Lilly has the lowest median risk rate. Such variation in risk rates between companies might reflect differences in their research areas as some diseases or targets can have systematically higher failure rate. Therefore, a high-risk pipeline portfolio might not be a reason to blame a company, but instead could reflect the fact that it searches for a treatment of a condition which is in general more complicated to treat. Nevertheless, our predictions can be of use in estimating the market risks of certain companies, especially small ones.

Currently, there is substantial pressure in the medical community to report all clinical trial results, and such transparency initiatives like AllTrials (www.alltrials.net) are gaining momentum (Powell-Smith and Goldacre 2016). Development of clinical trial prediction instruments and proof-of-concept demonstration may encourage big pharma companies to reveal some of the unreported clinical trial results and pay more attention to ML prediction methods.

## Conflict of Interest

All of the authors are employed by Insilico Medicine, Inc at the Pharmaceutical Artificial Intelligence (Pharma.AI) division, which is usually contracted by the large pharmaceutical companies to integrate deep learning solutions into the business practices. The publication of this paper will serve as an advertisement for some of these services. All of the authors declare conflict of interest.

## Acknowledgments

The authors would like to thank Nvidia for providing us with the TESLA K80 GPUs and for including us into the Inception program providing early access to materials and equipment. The authors cordially thank Polina Mamoshina and Franco Cortese of the Biogerontology Research Foundation for their valuable comments and editing of the manuscript.

## Supporting Information

**Figure S1.**
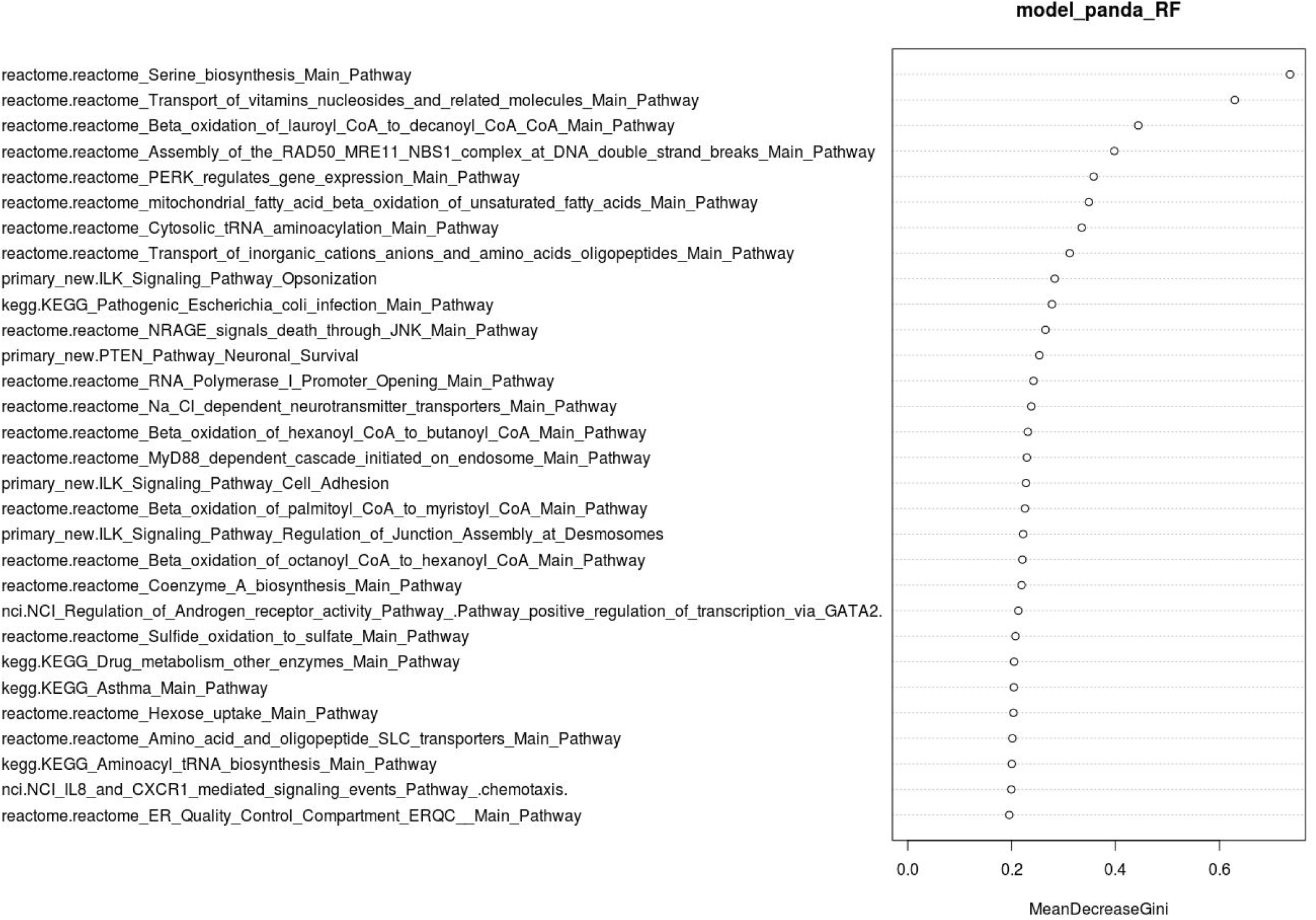
Variable importance of predictors for iPANDA pathway activation-based classifier. Notice the abundance of metabolic pathways (particularly, related to oxidation) and ER-related pathways.

## Supplementary text 1. Analysis of current R&D pipelines

In this section, we provide a succinct description of the current status of the development pipelines for five major pharmacology companies. The compounds described are currently at various stage of development, sometimes for many different indications related to oncology and cancer research. In some cases, they have received FDA approval and are already on the market. The information summarized below has been obtained from different sources including http://adisinsight.springer.com/, http://www.roche.com/research_and_development/who_we_are_how_we_work/pipeline.htm, http://www.merck.com/research/pipeline/home.html, http://www.abbvie.com/research-innovation/pipeline.html, https://www.lilly.com/pipeline/11.html, https://www.astrazeneca.com/our-science/pipeline.html

### Roche

Compound Alecensa (Alectinib), an anaplastic lymphoma kinase (ALK) inhibitor, is currently marketed for treatment against Non small cell lung cancer (NSCLC). Several Compounds are investigated regarding effects against diabetes mellitus type 2. This includes compound RO6870952 (target: GLP-1/GIP), currently in phase I, and Xenical (Orlistat) (target: pancreatic lipases) that has completed phase III and marketed. However, phase III has been discontinued for Aleglitazar (target: peroxisome proliferator-activated receptor) regarding its potential effects on diabetes mellitus type 2 and cardiovascular disorders. Boniva (target: osteoclast-mediated bone resorption) is marketed for treatment of cancer metastases but development was discontinued for treatment of cancer pain. Perjeta (Pertuzumab) (target: extracellular dimerization domain (Subdomain II) of the human epidermal growth factor receptor 2 protein (HER2)) is marketed for treatment of Breast cancer. It is in phase III for treatment of gastric cancer and ovarian cancer and in phase II for treatment of colorectal cancer. Its Development was discontinued for treatment of prostate cancer and NSCLC. Xeloda (Capecitabine), a thymidylate synthase inhibitor, is in phase II for treatment of glioma, oesophageal and pancreatic cancer. It is registered for treatment of rectal cancer and marketed for use against breast cancer, colorectal cancer and gastric cancer. However Xeloda was discontinued for treatment of Biliary cancer. Regarding Alzheimer's disease, Gantenerumab (target: Abeta fibrils) is in phase III, whereas Sembragiline (target: MAOB) successfully completed phase II. The development of MEM3454 (RG 3487) (target: nicotinic alpha7 receptor) has been discontinued. Gazyva (Obinutuzumab) is a CD20 antigen inhibitor and Immunomodulator. It is currently in phase II for treatment of follicular lymphoma and mantle-cell lymphoma and in phase III for diffuse large B cell lymphoma. Furthermore It is marketed for treatment of chronic lymphocytic leukaemia and non-Hodgkin's lymphoma. Inclacumab (target: P-selectin) was discontinued for treatment of acute coronary syndromes, myocardial infarction and peripheral vascular disorders. Lebrikizumab (target: IL-13) is in phase II for treatment of atopic dermatitis, chronic obstructive pulmonary disease and idiopathic pulmonary fibrosis and in phase III for treatment of asthma. It was discontinued for treatment of Hodgkin's disease. Zelboraf (Vemurafenib) is a proto oncogene protein b-raf inhibitor. It is in phase II for treatment of brain metastases, colorectal cancer, haematological malignancies, multiple myeloma, solid tumours and thyroid cancer. It is marketed for treatment of Malignant melanoma. Another successful compound is Mircera, an erythropoietin receptor agonist, used to treat patients with symptomatic anaemia associated with chronic kidney disease. One can also mention ORY-1001, a lysine specific demethylase 1 inhibitor, which is in phase I/II for treatment of acute myeloid leukaemia and in preclinical phase for treatment of acute lymphoblastic leukaemia and solid tumours and RG1662 (GABA A alpha 5 receptor modulator) in phase II for treatment of neurologic disorders.

### Merck

Several compounds are currently at different stages of development for various indications. MK-2206 (target: DPP-4) is in phase II for the treatment of breast cancer, colorectal cancer, haematological malignancies; NSCLC and solid tumours. Vorinostat (Histone deacetylase inhibitor) is in phase I for treatment against glioblastoma and Hodgkin's disease and in phase II for treatment of acute myeloid leukaemia, B cell lymphoma, breast cancer, glioma. It is marketed as a treatment against cutaneous T cell lymphoma. However, its development has been discontinued for treatment of gynaecological cancer, mesothelioma, multiple myeloma, non-Hodgkin's lymphoma, NSCLC and pancreatic cancer. Pembrolizumab (Keytruda) (CD274 antigen inhibitor; PDCD 1 protein inhibitor; programmed cell death 1 ligand 2 protein inhibitor) is marketed for treatment of head and neck cancer, malignant melanoma and NSCLC. It is pre-registered for use against Hodgkin's disease. Furthermore, it is in phase III for the treatment of breast cancer, colorectal cancer, gastric cancer, liver cancer, multiple myeloma, oesophageal cancer, renal cell carcinoma and urogenital cancer and in phase II for the treatment of adrenocortical carcinoma, bladder cancer, bone cancer, brain metastases, diffuse large B cell lymphoma, endometrial cancer, follicular lymphoma, glioblastoma, glioma, inflammatory breast cancer, merkel cell carcinoma, nasopharyngeal cancer, non-Hodgkin's lymphoma, ovarian cancer, pancreatic cancer, prostate cancer, rectal cancer, soft tissue sarcoma, solid tumours, thymoma and thyroid cancer. Finally, it is also in phase I/II for treatment of chronic lymphocytic leukaemia. Dalotuzumab (target: IGFR1) is in phase I/II for treatment of pancreatic cancer. It is also in phase II for treatment of breast cancer and colorectal cancer. Temodar (Temozolomide) (Alkylating agent; DNA cross linking agent; DNA synthesis inhibitor) is marketed as a treatment for anaplastic astrocytoma, glioma and malignant melanoma, it is in phase II for treatment of acute myeloid leukaemia and it has been discontinued for treatment of non-Hodgkin's lymphoma, prostate cancer and Sarcoma. Acadesine (Apoptosis stimulant; Platelet aggregation inhibitor; Purinergic P1 receptor agonist) has been discontinued for treatment of asthma, diabetes mellitus, ischaemic heart disorders and thrombosis. Anacetrapib (Cholesterol ester transfer protein inhibitor) is in phase III for treatment of atherosclerosis; hypercholesterolaemia; hyperlipidaemia; hyperlipoproteinaemia type IIa. Compound MK-1775 (AZD 1775) (WEE1 protein inhibitor) is in phase II for treatment of gynaecological cancer, NSCLC, ovarian cancer and SCLC. It is in phase I/II for treatment of solid tumours and in phase I for treatment of haematological disorders; head and neck cancer. However, it was discontinued for treatment of cervical cancer. Arcoxia (Etoricoxib) (Cyclo-oxygenase 2 inhibitor) is marketed for treatment of acute pain, ankylosing spondylitis, back pain, dental pain, dysmenorrhoea, gouty arthritis, musculoskeletal pain, osteoarthritis, postoperative pain, rheumatic disorders and rheumatoid arthritis. Finally one can mention the following compounds that are studied for one indication only. Compounds AF-130 (Purinergic P2X3 receptor antagonists) and ALX-0761 (Immunomodulator; Interleukin-17 inhibitor) are in phase I for action related to hypertension and migraine and psoriasis respectively. Atacicept (B cell activating factor inhibitor; Tumour necrosis factor ligand superfamily member 13 inhibitor) is in phase II/III for treatment of systemic lupus erythematosus and it has been discontinued for treatment of chronic lymphocytic leukaemia, lupus nephritis, multiple myeloma, multiple sclerosis, non-Hodgkin's lymphoma and rheumatoid arthritis. Odanacatib (Bone resorption factor inhibitor; Cathepsin K inhibitor) is in phase I for treatment of corticosteroid-induced osteoporosis but has been discontinued for various indications including bone metastases, male osteoporosis, osteoarthritis, osteoporosis and postmenopausal osteoporosis. Omarigliptin (CD26 antigen inhibitor) is marketed for treatment of diabetes mellitus Type 2 and Verubecestat (BACE1 protein inhibitor) is in phase III for treatment of Alzheimer's disease respectively. However three compounds, all targeting DPP-4, have seen their development discontinued. This includes Preladenant (Adenosine A2 receptor antagonist) for its potential action against Parkinson's disease, Vioxx (Rofecoxib), a Cyclo-oxygenase 2 inhibitor, for treatment of Alzheimer's disease, colorectal cancer; familial adenomatous polyposis, juvenile rheumatoid arthritis, Migraine, Periarthritis, sporadic adenomas. Furthermore, this component has been withdrawn from market for treatment of dental pain, dysmenorrhoea, musculoskeletal disorders, osteoarthritis, pain, rheumatoid arthritis. Tredaptive (GPR109A receptor agonist; GPR109B receptor agonist; Prostaglandin D2 receptor antagonist) has been withdrawn from market for treatment of dyslipidaemias, and hypercholesterolaemia and its development was discontinued for treatment of Atherosclerosis. Gardasil ( Immunostimulant) is marketed for treatment of anal cancer, anal intraepithelial neoplasia, cervical cancer, gynaecological cancer and is in phase III for treatment of Penile cancer. Januvia (Sitagliptin), a CD26 antigen inhibitor, is marketed for treatment of Type 2 diabetes mellitus but discontinued for Type 1 diabetes mellitus. PEG-Intron (Peginterferon alfa-2b) (Immunostimulant, Interferon alpha stimulants) is marketed for treatment of hepatitis B, hepatitis C and malignant melanoma. It has been discontinued for treatment of chronic myeloid leukaemia, multiple sclerosis, polycythaemia vera and solid tumours.

### Abbvie

Imbruvica (ibrutinib), a small molecule acting as a Bruton’s tyrosine kinase (BTK) inhibitor, is in phase II for the treatment of solid tumors, Multiple Myeloma (MM), Acute Myeloid Leukemia (AML) and Diffuse Large B-cell Lymphoma (DLBCL) and it is in phase III for chronic lymphocytic leukemia (CLL) and mantle cell lymphoma (MCL). Veliparib is a small molecule acting as an oral inhibitor targeting poly (adenosine diphosphate [ADP]-ribose) polymerase (PARP-1/2). It is being examined in combination with DNA-damaging therapies including chemotherapy or radiation. It is in phase I for its effects against solid tumors and non-small cell lung cancer (SCLC). It is also in phase II studies for colorectal cancer and in phase III for the treatment of ovarian cancer, triple-negative breast cancer, and HER2-negative metastatic or locally advanced BRCA breast cancer. Venclexta (venetoclax) is an oral B-cell lymphoma-2 (BCL-2) inhibitor which is in phase II for treatment of AML, DLBCL, non-Hodgkin lymphoma (NHL) and in phase III for the treatment of multiple myeloma (MM) and CLL. Xinlay (atrasentan) (target: Endothelin A receptor) is in phase III for the treatment of diabetic kidney disease and has completed phase III related to its effects on prostate cancer. ABT-767 is another PARP inhibitor currently under investigation and it is in phase I for treatment of ovarian cancer and various solid tumors. PCI-24781 is an inhibitor of histone deacetylase and has completed phase I for its effects against solid tumors. ABBV-8E12 (target: tau) is in phase I for treatment of neurologic disorders and a phase II is currently ongoing to evaluate the efficacy and safety of ABBV-8E12 in individuals with early Alzheimer’s disease (NCT02880956). ABT-414 is a monoclonal antibody drug conjugate targeting epidermal growth factor receptor (EGFR). It is a late stage investigational compound currently being studied in Phase I/II for the treatment of brain cancer and glioblastoma (GBM) whereas Rova-T (rovalpituzumab tesirine) is an antibody drug conjugate targeting the cancer stem cell-associated target delta-like protein 3 (DLL3). It is in phase I for the treatment of solid tumors and in phase II for treatment of SCLC and could be the first targeted therapy to show efficacy for this disease. PCI-27483, a factor VIIa inhibitors, was discontinued after Phase II trials for treatment of pancreatic cancer.

Finally, the following compounds have been approved by FDA: Venclexta is approved in the U.S. and indicated for the treatment of patients with relapsed/refractory CLL,Imbruvica is approved to treat patients with CLL and small lymphocytic lymphoma (SLL), Duodopa (target: hormonal system) is approved as a treatment against Parkinson's disease, and Elotuzumab (target: SLAMF7) has been approved for the treatment of multiple myeloma.

### Eli Lilly

Cyramza (ramucirumab) (target: VEGFR2) has completed phase II for treatment of prostate cancer, melanoma, ovarian cancer and renal cell carcinoma. It is also in phase III for treatment of Bladder cancer, hepatocellular carcinoma and gastric cancer. It is also in phase III studies for the treatment of first-line, EGFR positive NSCLC. Erbitux, an angiogenesis and epidermal growth factor (EGF) inhibitor, is approved and marketed for treatment of colorectal cancer, head and neck cancer. It has completed phase II for treatment of breast cancer. It is also in phase II for treatment of Bladder cancer, gastric cancer and rectal cancer. It is in phase III for treatment of non-small cell lung cancer. Gemzar (Gemcitabine), an inhibitor of DNA synthesis, is in phase I for treatment of head and neck cancer, in phase II for treatment of lymphoma and renal cell carcinoma and it has completed phase III for treatment against cervical dysplasia/cancer. Olaratumab (Platelet derived growth factor (PDGF) alpha receptor antagonist) is in phase I for treatment against solid tumours, it has completed phase II for its use against brain and prostate cancer. It is also in phase III for treatment of advanced sarcoma. Alimta (Pemetrexed) (target: cancer cell replication) is marketed for the treatment of mesothelioma and NSCLC. It has completed phase II for its use against breast cancer and it has also completed phase III for treatment against head and neck cancer but no further development was reported. Alimta was discontinued for the treatment of SCLC. Necitumumab (target: EGFR) is marketed as a treatment for NSCLC and has completed phase II for treatment of solid tumors. It was also in clinical trial for potential against colorectal cancer, but no development was reported for this indication. Abemaciclib, an inhibitor targeting Cyclin-dependent kinase 4 (CDK4) and Cyclin-dependent kinase 6 (CDK6) undergoes preclinical studies for treatment of solid tumors; it is in phase II for treatment of Cancer, Liposarcoma and Mantle-cell lymphoma. It is also in phase III for use against Non-small lung cancer and metastatic breast cancers. Gataparsen, a BIRC5 protein and protein synthesis inhibitor, was discontinued for treatment of acute myeloid leukaemia and has also been discontinued for treatment of NSCLC and prostate cancer after completion of phase II. Compounds that are under study for one indication are also worth mentioning. Tabalumab, an inhibitor targeting B cell activating factor inhibitors (BAF) has been discontinued after phase II/III for treatment of multiple myeloma, Taltz, an Interleukin-17 inhibitors, has completed phase II for its action against rheumatoid arthritis. Teplizumab, CD3 antigen inhibitors, is in phase II for type 1 diabetes mellitus. Galunisertib is a chemical compound that blocks transforming growth factor-*β* receptor I kinase and TGF-*β* signaling. It is currently under phase II for the treatment of hepatocellular cancer. Merestinib is another chemical compound that has been shown to be a reversible type II ATP-competitive inhibitor of MET a receptor for hepatocyte growth factor and is currently in phase II for the treatment of cancer. Prexasertib (checkpoint kinase 1 (Chk1) inhibitor) is also in phase II for the treatment of cancer as well as Ralimetinib (p38 MAP kinase inhibitor) in phase II for treatment of ovarian cancer. Chemical entities currently in phase I for the treatment of cancer include Angiopoietin 2 antibody, Chk1 Inhibitor, CSF-1R Monoclonal Antibody, ERK inhibitor, FGFR3-ADC and PD-L1 Antibody. However, for several compounds, clinical studies have been unsuccessful for some indications. Zosuquidar, a P-glycoprotein inhibitors, was discontinued for treatment of acute myeloid leukaemia after completion of phases I/II and III. One can mention Arzoxifene and Evacetrapib that have been discontinued regarding their potential use against breast cancer and cardiovascular disorders. Erbitux was discontinued regarding its potential actions against lung cancer. Enzastaurin (target: PKC) has been discontinued for treatment of brain and lung cancer. Peglispro (target: glucose uptake) has been discontinued regarding its potential action against diabetes mellitus type 1 and type 2.

Other compounds have received FDA approval. This includes: Alimta is approved by FDA for the treatment of lung cancer. Basaglar (target: glucose uptake) has FDA approval for its use against diabetes mellitus type 1 and type 2. Necitumumab received FDA approval for its use in the treatment of lung cancer. Cyramza (target: VEGFR2) received FDA approval to be used against colorectal, gastric and lung cancers. Erbitux received FDA approval for its use against colorectal cancer and head and neck cancer. Cymbalta (target: hormonal system) for the treatment of pain, acute or chronic, Dulaglutide (target: GLP-1 receptor) and Jentadueto (target: DPP-4) for the treatment of diabetes mellitus type 2 and Raloxifene (target: estrogen receptors) for the treatment of breast cancer. Gemzar received FDA approval and is marketed for treatment of biliary cancer, Bladder cancer, breast cancer, NSCLC, ovarian cancer and pancreatic cancer.

### AstraZeneca

Recentin (Cediranib) (target: VEGFR) is in phase II/III for treatment of ovarian cancer and has been discontinued for various indications including acute myeloid leukaemia, biliary cancer, breast cancer, colorectal cancer, gastric cancer, gastrointestinal stromal tumours, glioblastoma; NSCLC, renal cell carcinoma, soft tissue sarcoma, solid tumours. Selumetinib ( MAP kinase kinase 1 and 2 inhibitor) is in phase III for treatment of thyroid cancer, in phase II for treatment of biliary cancer, colorectal cancer, neurofibromatoses and NSCLC. It is also in phase I/II for treatment of astrocytoma and glioma, in phase I for treatment of solid tumours and in preclinical stage for treatment of pancreatic cancer. It was discontinued for the treatment of acute myeloid leukaemia, hepatocellular carcinoma, malignant melanoma, multiple myeloma and uveal melanoma. Lynparza ( Olaparib), a Poly(ADP-ribose) polymerase inhibitor, is marketed for the treatment of ovarian cancer, it is in phase III for treatment of breast cancer and pancreatic cancer, in phase II for treatment of prostate cancer and solid tumours and in Phase I/II for treatment of fallopian tube cancer, glioblastoma, head and neck cancer, peritoneal cancer and NSCLC. Iressa (Gefitinib) (Epidermal growth factor receptor antagonist) is marketed for the treatment of NSCLC, it is in phase II for treatment of salivary gland cancer and in phase I for the treatment of head and neck cancer. It was discontinued for the treatment of Bladder cancer, brain metastases, breast cancer, colorectal cancer, glioblastoma, renal cell carcinoma and thyroid cancer.

Compound AZD2423 (Chemokine inhibitor) was discontinued for the treatment of chronic obstructive pulmonary disease (COPD) and neuropathic pain. Brilinta (Ticagrelor) (Platelet ADP receptor antagonist and Purinoceptor P2Y12 antagonist) is marketed for the treatment of acute coronary syndromes, it is in phase III for the treatment of cardiovascular disorders and stroke and in phase II for the treatment of abdominal aortic aneurysm, community-acquired pneumonia, coronary artery disease, sickle cell anaemia and it is has been discontinued for the treatment of peripheral arterial disorders. Crestor (Rosuvastatin) (HMG-CoA reductase inhibitor) is marketed for the treatment of atherosclerosis, cardiovascular disorders, hypercholesterolaemia, hyperlipidaemia, hypertriglyceridaemia and was discontinued for the treatment of Heart failure. AZD2811 (Aurora kinase B inhibitor) is in phase I for treatment against solid tumors and in preclinical stage for the treatment of diffuse large B cell lymphoma and lung cancer. Tagrisso (Osimertinib) (Epidermal growth factor receptor antagonist and protein tyrosine kinase inhibitor) is marketed for the treatment of NSCLC, in phase II for the treatment of carcinomatous meningitis and in phase I for the treatment of solid tumours. Anastrozole (Aromatase inhibitor and estrogen receptor antagonists)and Faslodex (Fulvestrant) (Estrogen receptor antagonist and selective estrogen receptor degrader) are marketed for treatment of breast cancer. Casodex (Bicalutamide) (Testosterone congener inhibitor) is marketed for the treatment of prostate cancer. Daliresp (Roflumilast) (Type 4 cyclic nucleotide phosphodiesterase inhibitor), Symbicort (Budesonide/formoterol) (Beta 2 adrenergic receptor agonist, Immunosuppressant and Steroid receptor agonist) and Bevespi (Formoterol/glycopyrrolate) (Beta 2 adrenergic receptor agonist and muscarinic receptor antagonist) are marketed for the treatment of COPD. Tomudex (Raltitrexed) (Thymidylate synthase inhibitor) is marketed for the treatment of colorectal cancer and mesothelioma.

## References

Aliper, Alexander, Sergey Plis, Artem Artemov, Alvaro Ulloa, Polina Mamoshina, and Alex Zhavoronkov. 2016. “Deep Learning Applications for Predicting Pharmacological Properties of Drugs and Drug Repurposing Using Transcriptomic Data.” Molecular Pharmaceutics 13 (7): 2524–30.

Ashburn, Ted T., and Karl B. Thor. 2004. “Drug Repositioning: Identifying and Developing New Uses for Existing Drugs.” Nature Reviews. Drug Discovery 3 (8): 673–83.

Berry, Donald A. 2011. “Adaptive Clinical Trials in Oncology.” Nature Reviews. Clinical Oncology 9 (4): 199–207.

Bickerton, G. Richard, Gaia V. Paolini, Jérémy Besnard, Sorel Muresan, and Andrew L. Hopkins. 2012. “Quantifying the Chemical Beauty of Drugs.” Nature Chemistry 4 (2): 90–98.

Brown, Elliot G., Louise Wood, and Sue Wood. 1999. “The Medical Dictionary for Regulatory Activities (MedDRA).” Drug Safety: An International Journal of Medical Toxicology and Drug Experience 20 (2): 109–17.

Chabanas, Matthieu, Vincent Luboz, and Yohan Payan. 2003. “Patient Specific Finite Element Model of the Face Soft Tissues for Computer-Assisted Maxillofacial Surgery.” Medical Image Analysis 7 (2): 131–51.

Clermont, Gilles, John Bartels, Rukmini Kumar, Greg Constantine, Yoram Vodovotz, and Carson Chow. 2004. “In Silico Design of Clinical Trials: A Method Coming of Age.” Critical Care Medicine 32 (10): 2061–70.

Cohen, Alan A., Vincent Morissette-Thomas, Luigi Ferrucci, and Linda P. Fried. 2016. “Deep Biomarkers of Aging Are Population-Dependent.” Aging 8 (9): 2253–55.

DiMasi, Joseph A., Henry G. Grabowski, and Ronald W. Hansen. 2016. “Innovation in the Pharmaceutical Industry: New Estimates of R&D Costs.” Journal of Health Economics 47: 20–33.

Duchi, John, Elad Hazan, and Yoram Singer. 2011. “Adaptive Subgradient Methods for Online Learning and Stochastic Optimization.” Journal of Machine Learning Research: JMLR, July.

Evens, Andrew M., Sriram Balasubramanian, Julie M. Vose, Wael Harb, Leo I. Gordon, Robert Langdon, Julian Sprague, et al. 2016. “A Phase I/II Multicenter, Open-Label Study of the Oral Histone Deacetylase Inhibitor Abexinostat in Relapsed/Refractory Lymphoma.” Clinical Cancer Research: An Official Journal of the American Association for Cancer Research 22 (5): 1059–66.

Fernandez, J. W., and P. J. Hunter. 2005. “An Anatomically Based Patient-Specific Finite Element Model of Patella Articulation: Towards a Diagnostic Tool.” Biomechanics and Modeling in Mechanobiology 4 (1): 20–38.

Gayvert, Kaitlyn M., Neel S. Madhukar, and Olivier Elemento. 2016. “A Data-Driven Approach to Predicting Successes and Failures of Clinical Trials.” Cell Chemical Biology 23 (10): 1294–1301.

Ghose, A. K., V. N. Viswanadhan, and J. J. Wendoloski. 1999. “A Knowledge-Based Approach in Designing Combinatorial or Medicinal Chemistry Libraries for Drug Discovery. 1. A Qualitative and Quantitative Characterization of Known Drug Databases.” Journal of Combinatorial Chemistry 1 (1): 55–68.

Goodfellow, Ian, Jean Pouget-Abadie, Mehdi Mirza, Bing Xu, David Warde-Farley, Sherjil Ozair, Aaron Courville, and Yoshua Bengio. 2014. “Generative Adversarial Nets.” Advances in Neural Information Processing Systems, 2672–80.

Hay, Michael, David W. Thomas, John L. Craighead, Celia Economides, and Jesse Rosenthal. 2014. “Clinical Development Success Rates for Investigational Drugs.” Nature Biotechnology 32 (1): 40–51.

Jeliazkova, Nina, and Vedrin Jeliazkov. 2011. “AMBIT RESTful Web Services: An Implementation of the OpenTox Application Programming Interface.” Journal of Cheminformatics 3 (May): 18.

Kadurin, Artur, Alexander Aliper, Andrey Kazennov, Polina Mamoshina, Quentin Vanhaelen, Kuzma Khrabrov, and Alex Zhavoronkov. 2016. “The Cornucopia of Meaningful Leads: Applying Deep Adversarial Autoencoders for New Molecule Development in Oncology.” Oncotarget, December. doi:10.18632/oncotarget.14073.

Kanehisa, M. 2000. “KEGG: Kyoto Encyclopedia of Genes and Genomes.” Nucleic Acids Research 28 (1): 27–30.

Kingma, Diederik, and Jimmy Ba. 2014. “Adam: A Method for Stochastic Optimization.” arXiv.

Kuhn, Michael, Ivica Letunic, Lars Juhl Jensen, and Peer Bork. 2015. “The SIDER Database of Drugs and Side Effects.” Nucleic Acids Research 44 (D1): D1075–79.

Lamb, Justin, Emily D. Crawford, David Peck, Joshua W. Modell, Irene C. Blat, Matthew J. Wrobel, Jim Lerner, et al. 2006. “The Connectivity Map: Using Gene-Expression Signatures to Connect Small Molecules, Genes, and Disease.” Science 313 (5795): 1929–35.

Lavine, Kory J., and Douglas L. Mann. 2013. “Rethinking Phase II Clinical Trial Design in Heart Failure.” Clinical Investigation 3 (1): 57–68.

LeCun, Yann, Yoshua Bengio, and Geoffrey Hinton. 2015. “Deep Learning.” Nature 521 (7553): 436–44.

Ledford, Heidi. 2011. “Translational Research: 4 Ways to Fix the Clinical Trial.” Nature 477 (7366): 526–28.

Leeson, Paul D., and Brian Springthorpe. 2007. “The Influence of Drug-like Concepts on Decision-Making in Medicinal Chemistry.” Nature Reviews. Drug Discovery 6 (11): 881–90.

Li, Nicole Y. K., Katherine Verdolini, Gilles Clermont, Qi Mi, Elaine N. Rubinstein, Patricia A. Hebda, and Yoram Vodovotz. 2008. “A Patient-Specific in Silico Model of Inflammation and Healing Tested in Acute Vocal Fold Injury.” PloS One 3 (7): e2789.

Lipinski, C. A., F. Lombardo, B. W. Dominy, and P. J. Feeney. 2001. “Experimental and Computational Approaches to Estimate Solubility and Permeability in Drug Discovery and Development Settings.” Advanced Drug Delivery Reviews 46 (1-3): 3–26.

Makhzani, Alireza, Jonathon Shlens, Navdeep Jaitly, Ian Goodfellow, and Brendan Frey. 2015. “Adversarial Autoencoders.” bioRxiv.

Mamoshina, Polina, Armando Vieira, Evgeny Putin, and Alex Zhavoronkov. 2016. “Applications of Deep Learning in Biomedicine.” Molecular Pharmaceutics 13 (5): 1445–54.

Mignani, Serge, Scot Huber, Helena Tomás, João Rodrigues, and Jean-Pierre Majoral. 2016. “Why and How Have Drug Discovery Strategies in Pharma Changed? What Are the New Mindsets?” Drug Discovery Today 21 (2): 239–49.

Mita, Monica M., Anil A. Joy, Alain Mita, Kamalesh Sankhala, Ying-Ming Jou, Da Zhang, Paul Statkevich, et al. 2014. “Randomized Phase II Trial of the Cyclin-Dependent Kinase Inhibitor Dinaciclib (MK-7965) versus Capecitabine in Patients with Advanced Breast Cancer.” Clinical Breast Cancer 14 (3): 169–76.

Mullard, Asher. 2013. “2012 FDA Drug Approvals.” Nature Reviews. Drug Discovery 12 (2): 87–90.

Munos, Bernard H., and William W. Chin. 2011. “How to Revive Breakthrough Innovation in the Pharmaceutical Industry.” Science Translational Medicine 3 (89): 89cm16.

Ozerov, Ivan V., Ksenia V. Lezhnina, Evgeny Izumchenko, Artem V. Artemov, Sergey Medintsev, Quentin Vanhaelen, Alexander Aliper, et al. 2016. “In Silico Pathway Activation Network Decomposition Analysis (iPANDA) as a Method for Biomarker Development.” Nature Communications 7 (November): 13427.

Patlewicz, Grace, and Jeremy M. Fitzpatrick. 2016. “Current and Future Perspectives on the Development, Evaluation, and Application of in Silico Approaches for Predicting Toxicity.” Chemical Research in Toxicology 29 (4): 438–51.

Powell-Smith, Anna, and Ben Goldacre. 2016. “The TrialsTracker: Automated Ongoing Monitoring of Failure to Share Clinical Trial Results by All Major Companies and Research Institutions.” F1000Research 5: 2629.

Putin, Evgeny, Polina Mamoshina, Alexander Aliper, Mikhail Korzinkin, Alexey Moskalev, Alexey Kolosov, Alexander Ostrovskiy, Charles Cantor, Jan Vijg, and Alex Zhavoronkov. 2016. “Deep Biomarkers of Human Aging: Application of Deep Neural Networks to Biomarker Development.” Aging 8 (5): 1021–33.

Rubin, Eric H., and D. Gary Gilliland. 2012. “Drug Development and Clinical Trials–the Path to an Approved Cancer Drug.” Nature Reviews. Clinical Oncology 9 (4): 215–22.

Scannell, Jack W., Alex Blanckley, Helen Boldon, and Brian Warrington. 2012. “Diagnosing the Decline in Pharmaceutical R&D Efficiency.” Nature Reviews. Drug Discovery 11 (3): 191–200.

Schaefer, Carl F., Kira Anthony, Shiva Krupa, Jeffrey Buchoff, Matthew Day, Timo Hannay, and Kenneth H. Buetow. 2009. “PID: The Pathway Interaction Database.” Nucleic Acids Research 37 (Database issue): D674–79.

Srivastava, Nitish, Geoffrey Hinton, Alex Krizhevsky, Ilya Sutskever, and Ruslan Salakhutdinov. 2014. “Dropout: A Simple Way to Prevent Neural Networks from Overfitting.” Journal of Machine Learning Research: JMLR, June, 1929–1958.

Stephenson, Joe J., John Nemunaitis, Anil A. Joy, Julie C. Martin, Ying-Ming Jou, Da Zhang, Paul Statkevich, et al. 2014. “Randomized Phase 2 Study of the Cyclin-Dependent Kinase Inhibitor Dinaciclib (MK-7965) versus Erlotinib in Patients with Non-Small Cell Lung Cancer.” Lung Cancer 83 (2): 219–23.

Vanhaelen, Quentin, Polina Mamoshina, Alexander M. Aliper, Artem Artemov, Ksenia Lezhnina, Ivan Ozerov, Ivan Labat, and Alex Zhavoronkov. 2016. “Design of Efficient Computational Workflows for in Silico Drug Repurposing.” Drug Discovery Today, September. doi:10.1016/j.drudis.2016.09.019.

Veber, Daniel F., Stephen R. Johnson, Hung-Yuan Cheng, Brian R. Smith, Keith W. Ward, and Kenneth D. Kopple. 2002. “Molecular Properties That Influence the Oral Bioavailability of Drug Candidates.” Journal of Medicinal Chemistry 45 (12): 2615–23.

Viceconti, Marco, Adriano Henney, and Edwin Morley-Fletcher. 2016. “In Silico Clinical Trials: How Computer Simulation Will Transform the Biomedical Industry.” International Journal of Clinical Trials 3 (2): 37.

Zeiler, Matthew D. 2012. “ADADELTA: An Adaptive Learning Rate Method.”

